# Stochasticity and Bet Hedging Drive Cryptococcal Capsule Dynamics

**DOI:** 10.64898/2026.06.17.732865

**Authors:** Quigly Dragotakes, Lia Sanchez Ramirez, Arturo Casadevall

## Abstract

*Cryptococcus neoformans* and related species are major human pathogens that cause cryptococcosis, a disease with high mortality and morbidity despite antifungal therapy. Pathogenic *Cryptococcus* spp. cells express a polysaccharide capsule, which is the most important virulence factor. In this study we analyzed the distribution of capsule sizes for several strains from *Cryptococcus spp*. and found that they follow stochastic dynamics, with a heavy right-hand tail distribution, favoring larger capsules. The distribution for each strain is remarkably stable despite repeated perturbation of culture conditions including media refreshment, time, and macrophage ingestion. Growth in macrophages resulted in different capsule distributions, observed in vitro, with a suggestion of different polysaccharide-like materials formed or utilized in the resident phagosome. We propose that the stability in capsule size distributions represents a ‘capsulestat’ mechanism for the population. An emergent property whereby individual cells manifest capsule size variation emanating from random effects on individual capsule assembly steps. This distribution balances between cells with large capsules that are less susceptible to a variety of environmental stresses at the price of slower replication, increased size, and increased energy requirements and cells with smaller, less protective capsules that reproduce faster. Thus, *Cryptococcus spp*. populations establish a bet hedging strategy that can enhance the viability of the population as conditions change at the cost of optimal short-term growth.

## Introduction

*Cryptococcus neoformans* is a pathogenic yeast responsible for hundreds of thousands of deaths annually, primarily in immune compromised populations. Conversely, *Cryptococcus gattii* affects fewer individuals but is capable of dissemination in immune competent hosts^1^. The pathogenic success of these *Cryptococcus spp*. in mammalian hosts is due to several virulence factors which allow the yeast to survive the phagolysosome and evade immune defenses^2^.

One of the most important virulence factors of pathogenic *Cryptococcus* spp. is the polysaccharide capsule^3^. This capsule protects the yeast from desiccation, environmental acidity, ROS stress, UV stress, antibody binding, and phagocytosis. The capsule also protects cryptococcal cells against their primary environmental predators: amoeba^4^. Multivariate linear regression of the contribution of the various cryptococcal virulence factors to virulence shows that the capsule makes the largest contribution to its virulence^5^. Consequently, understanding the dynamics behind capsule growth is imperative to understanding cryptococcal pathogenesis. This capsule is constructed of three major components: glucuronoxylomannan (GXM), galactoxylomannan (GalXM), and mannoproteins. Capsule growth is induced under various stress conditions and protects the yeast from a wide array of cellular stresses^2,6^. Specifically, this robust capsule offers protection from many of the stresses present in the phagolysosome including acidity and reactive oxygen species (ROS)^7^. We have previously shown that macrophages hedge bets during phagosome acidification, optimizing their ability to inhibit ingested pathogens^8^. Thus, we hypothesized that the sizes of cryptococcal capsules are similarly distributed to combat predation by different environmental phagocytic predators and, due to similarities, the macrophage betting strategy.xs

In this study, we investigated the dynamics of capsule growth for *C. neoformans* and *C. gattii* strains in a variety of growth conditions to determine how the capsule is grown, what distribution of capsule size yeast populations achieve, and what benefits are associated with the specific capsule size distribution we observe. Previous work in the field has established general benefits to increased capsule size at the population level, which we expand upon here by increasing resolution to individual capsule size and comparing those benefits to an estimated cost of capsule synthesis.

## Methods

### Yeast Strains and Culture Conditions

*C. neoformans* strain H99 and *C. gattii* strains WM179, R265, and WM1611 were cultured from frozen stocks on yeast potato dextrose (YPD) agar plates, then seeded into 3 mL YPD broth for 2 d before use in experiments. An *C. neoformans* H99 *actin:GFP* strain was sourced from the May lab^9^. Capsule induction was achieved by culturing cells in minimal media (MM): 15 mM dextrose, 10 mM MgSO_4_, 29.3 mM KH_2_PO_4_, 13 mM glycine, and 3 µM thymine-HCl at pH 5.5.

For long term capsule induction experiments, *C. neoformans* was cultured in 3 mL MM on a rotational culture wheel. Each week 50 µL was sampled to measure capsule size by optical microscopy via India Ink exclusion. If the sample was refreshed, the remaining culture was pelleted and resuspended in fresh MM. Otherwise, the sample was placed back on the wheel with no further modification.

### Mammalian Cell Culture and Infections

Bone marrow derived macrophages (BMDMs) were harvested from 4-6 week old C57BL/6J female mouse hind leg bones and differentiated over 7 d in BMDM media (10% FBS, 20% conditioned L929 supernatant, 1% HEPES, 1% Penicillin/streptomycin, 1% GlutaMAX, 1% Non-essential amino acids, and 0.1% betamercaptoethanol in DMEM base). Macrophages were then seeded at 10^6^ cells / well in 6-well tissue culture plates and polarized overnight: IFN-γ (10 ng / mL) and LPS (0.5 µg / mL) for M1 or IL-4 (20 ng / mL) for M2. BMDMs were refreshed with new media and infected with guinea pig complement opsonized or unopsonized yeast at an MOI of 1 for 24 h. BMDMs were harvested with non-enzymatic cellstripper and lysed by incubation in dH_2_O for 15 min followed by 10 syringe pulls through a 26G needle. Released intracellular yeast were then collected by centrifugation at 2300 *g* for 5 min.

### Capsule and Cell Body Measurements

India Ink slides were prepared by mixing 6 μL of cell suspension with 4 μL India Ink onto a glass slide. The slides were then imaged under brightfield microscopy (Olympus AX70 or LEICA Thunder) and capsules measured by India Ink exclusion zone^10^.

### Chaos Analysis

To investigate chaotic or stochastic signatures within data, we used the previously established scanning window analysis^8,11,12^. Briefly, intervals between measurements are arranged in a vector. A scanning window of 4 units is then used to assign each window an ordinal pattern from smallest to largest value within that window. The frequency of each ordinal pattern is then calculated. A stochastic system will show every pattern in some non-zero frequency, while a deterministic system will have certain “forbidden patterns” that never appear.

### ROS Susceptibility Assay

*C. neoformans* cultures were grown in YPD or MM for 5 d, then washed with PBS and counted via hemocytometer. The two cultures were normalized to equivalent cell counts in 500 µL total PBS by lowest common denominator, two samples for each culture condition were prepared. One sample from each culture condition was treated with 2 mM H_2_O_2_ for 1.5 h. The cells were then imaged via bright field microscopy (Olympus AX70) with India Ink and fluorescence at 488 nm / 530 nm. We then measured the size of each observed yeast and recorded whether it was GFP positive. GFP negative cells are considered inhibited. Viability statistics were calculated using the test of equal proportions.

### Growth Curves

*C. neoformans* cultures were seeded at 10^4^ cell / mL in 1.5 mL YPD per well in 12 well tissue culture plates. Cultures were then incubated for 48-72 h with orbital shaking at 23, 30, or 37 °C and absorbance read at 600 nm every 1-5 min.

### Estimating Capsule Energy Investiture and Sugar Consumption

Previous work has characterized the density of polysaccharide at various locations throughout the *C. neoformans* capsule with two main regions (high density and low density). For the sake of this model, we assume the capsule is made entirely of GXM while in reality, GXM is only estimated to make up 90% of the capsule. Thus, we use the following equation to calculate J / µm of capsule radius. First, because the density of GXM changes throughout the capsule, we must calculate the high density and low-density region energy requirements separately:

Energy in High Density Region (< 1.5 µm) + Energy in Low Density Region (> 1.5 µm)

Next, because these regions are hollow spheres, we subtracted from each region the radii of the cell body and cell body plus high-density capsule regions, respectively:

[Energy in High Density Region – Energy of region occupied by cell body] + [Energy of Low Density Region – Energy of region occupied by High Density Region and Cell Body]

Finally, we can calculate each of these regions with the general equation:

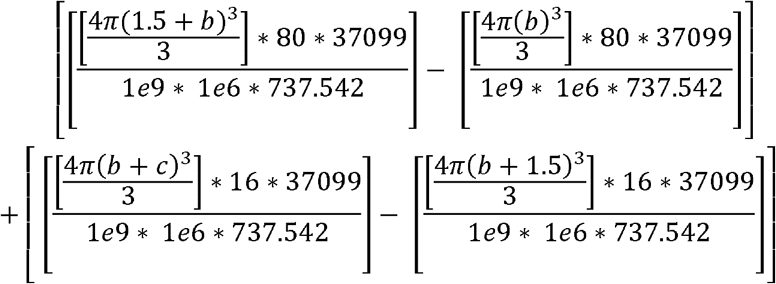

Wherein *c* is the radius of a given yeast cell capsule, *b* is the radius of a given yeast cell body, 16 µg / µL is the estimated density of capsular polysaccharide past 1.5 µm, 737.542 g / mol is the estimated molecular weight of serotype D GXM, and 37099 kJ / mol is the estimated enthalpy of serotype D GXM. Cells with a capsule radius smaller than 1.5 µm are filtered out logically and calculated only using the first half of the equation, since they would not have a region of low-density polysaccharide.

We will assume a population of *Cryptococcus neoformans* cells in a volume can occupy approximately 70% of the total free space since rigid spheres cannot pack perfectly into a volume and, under usual circumstances, we do not see capsules overlapping. We can then use the previous equation to predict what concentration of glucose would be required to sustain said population per cubic centimeter, assuming respiration produces approximately 500 kJ / mol O_2_.

It should be noted that these models rely on several assumptions and only represent a floor estimate of the energy involved. We assume perfect conversion of glucose to energy in respiration and did not account for energy required to export sugar monomers, construct those monomers into more complex capsular structures, or subunits aside from GXM serotype D.

## Results

### Cryptococcus neoformans Capsule Distribution Dynamics Change with Growth Conditions

We first investigated the capsule distribution of *C. neoformans* in standard growth conditions by culturing *C. neoformans* in either YPD for 2 d or minimal media for 5 d, then measuring the capsule size of individual cells. We observed differences in the two capsule size distributions with the *C. neoformans* strain H99 cells grown in YPD being closer to normality and those grown in MM being more variable with a heavy tail (Figure 1A, B).

**Figure 1:**
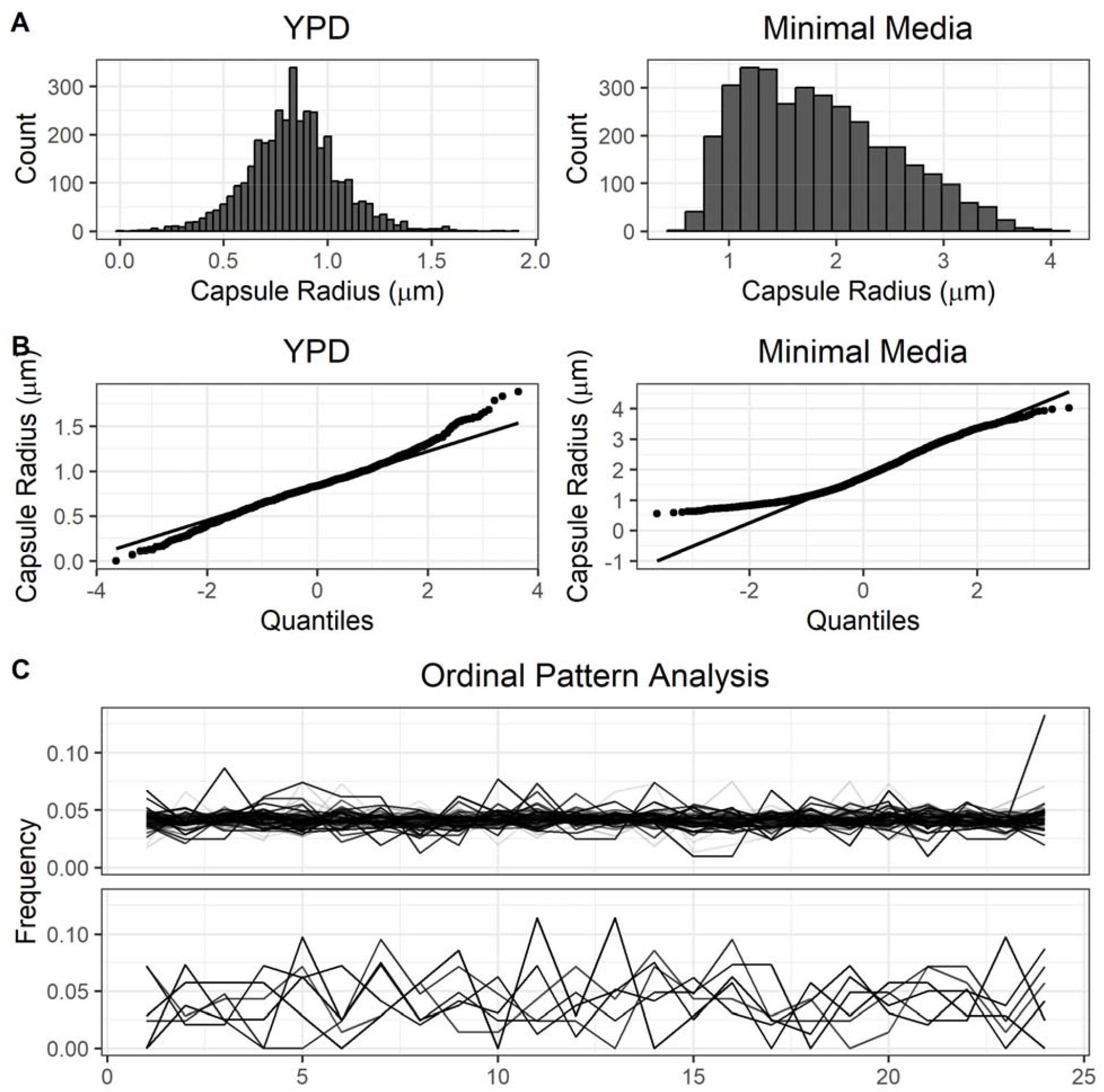
Observed capsule sizes of *C. neoformans* strain H99 grown under nutrient rich (YPD) or nutrient starved (Minimal Media) conditions. **A.** Distributions of capsule radii for each media condition (n > 3000) **B**. QQ plots of capsule radii for each media condition. Neither distribution satisfies normality tests, but the YPD grown distribution approaches normality much closer than the MM distribution. The MM distribution displays heavy tail skewing. **C**. Ordinal pattern analysis for every set of capsule radius data found in this paper. Only four conditions exhibited chaotic forbidden ordinals, however those six conditions lacked power due to reduced sample sizes to suggest the existence of chaos.

We next sought to determine the dynamics of capsule generation under these conditions by employing a scanning window ordinal pattern analysis to each experiment. Only six conditions exhibited chaotic signatures of forbidden ordinals, however these forbidden patterns are more likely due to reduced sample sizes in these conditions (n < 350 cells) than true chaos (Figure 1C) ^13^. Thus, *C. neoformans* final capsule size is likely determined randomly from a set distribution which changes depending on environmental conditions.

Interestingly, the capsule size distribution for cells grown in MM displayed significant skewing from a normal distribution. However, we were not sure if this skew was a legitimate reflection of the population, or if the cultures simply needed more time to increase the capsule size of the total population, eventually resulting in a normal distribution with a shifted average compared to YPD cultures.

### Cryptococcus neoformans Exhibits a Stable Distribution of Capsule Sizes

To investigate potential temporal differences in capsule dynamics, we measured capsule sizes over a three-month period under differing resource conditions. *C. neoformans* cells were cultured in MM either refreshed weekly or not refreshed. Half of the samples in each condition were switched after the first four weeks. Surprisingly, we found that the distribution of capsule sizes among all conditions over all weeks studied had only slight variations (Figure 2A). Furthermore, by the end of the first day we observed a distribution similar to that of the following time intervals, suggesting that the capsule is induced quite early on during growth in media and remains stable (Figure 2B). In fact, we were able to observe individual yeast with enlarged capsules as early as 1 h post incubation, though the population dynamics shifted closer to the 24 h mark. Overall, capsules at early times were relatively normally distributed and we did not see major capsule induction until about 16 h, at which point the distribution began to skew (Figure 2C). These data suggested our initial hypothesis was wrong and that increasing amounts of time in capsule inducing conditions will not result in a YPD-like normal distribution of capsule sizes.

**Figure 2:**
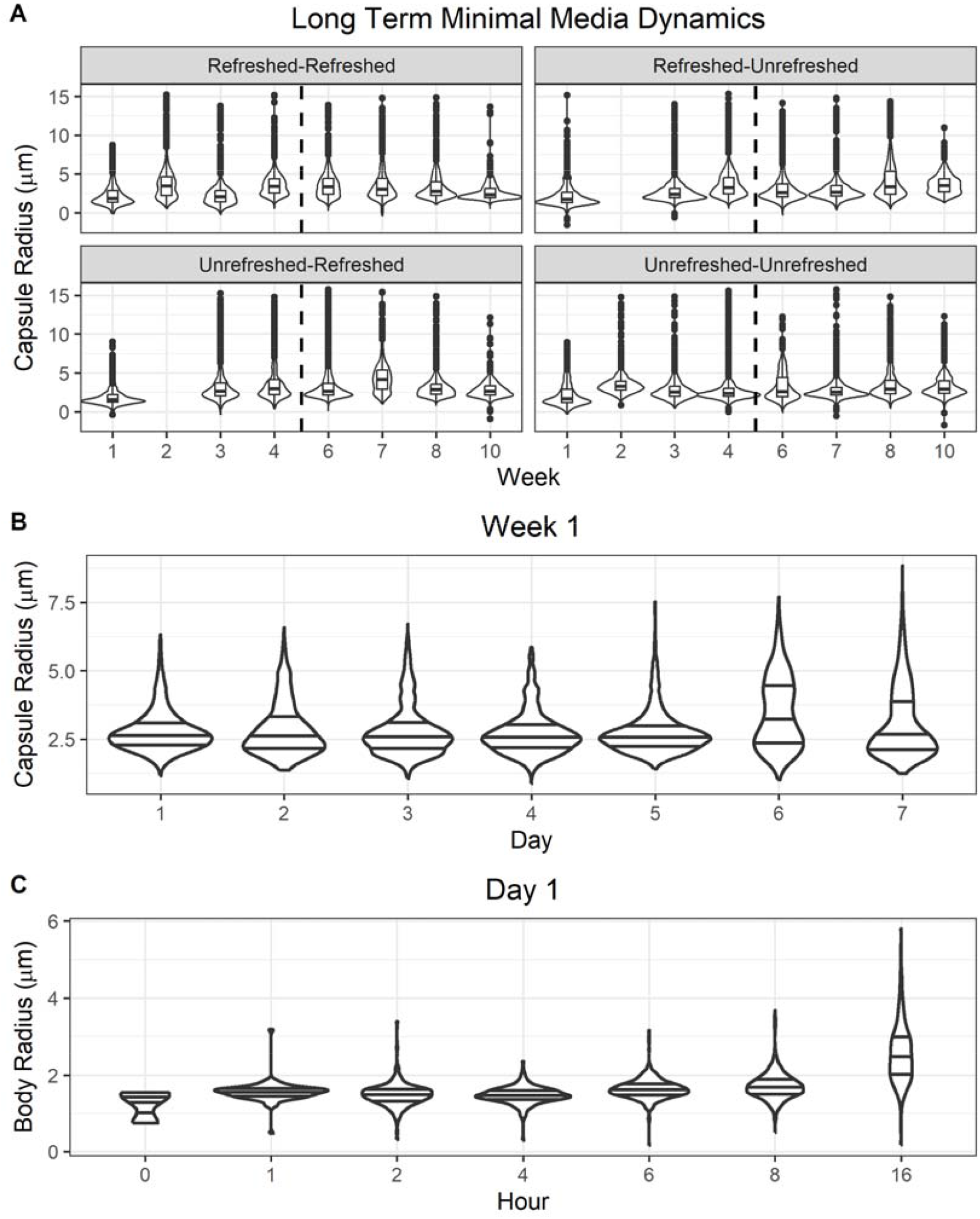
Capsule dynamics of *C. neoformans* strain H99 in minimal media with and without weekly refreshing of media. **A.** Week by week capsule measurements of cultures whose media was or was not refreshed after measurement each week. After week 4 (dotted line) the refreshed status of half the cultures were switched. Each of the four conditions is representative of three biological replicates. **B**. Day by day measurements of the first week of incubation. **C**. Hour by hour of the first day of incubation. Capsule distributions were surprisingly stable throughout all time frames. Every violin is a composite of at least three biological replicates and n > 550.

Given the remarkably stable capsule distributions we observed over 12 weeks of incubation and regardless of nutritional refreshment, we investigated whether this distribution was achieved at the population or individual level. Put simply, do *C. neoformans* yeast inherit a capsule size from their parent that remains relatively constant, or does each new yeast randomly determine capsule size. We observed consistent distributions between individual colony replicates with right tail skewing (Figure 2D). Observing that the same right hand skewed distribution arose from single colonies suggested that capsule size was not inherited, and that each new yeast randomly determines its own capsule size. In fact, the distribution of capsule sizes in capsule inducing conditions was remarkably consistent across all samples and experiments, always resembling a normal distribution but with a heavy right-hand tail (Supplemental Figure 1). We suspect the strong conservation of capsule distribution implies an inherent benefit to the population, and that this particular distribution arose due to selective pressures from the environment.

### Overall Capsule Distribution Stability is Conserved Within and Between Species

Given the known differences in virulence and our observed differences between *C. neoformans* and *C. gattii* capsule dynamics during macrophage infection, we sought to expand all experiments with three *C. gattii* strains as well as an additional *C. neoformans* strain KN99. Surprisingly, we found that capsule dynamics were highly conserved between species and strains with only slight differences compared to *C. neoformans* strain H99.

Each *C. gattii* strain had larger average capsule size compared to *C. neoformans* H99 but these also differed in each strain between intracellular and extracellular yeast (Figure 4D). Most notably, *C. gattii* strain WM161 had a particularly large and normally distributed population of extracellular capsule sizes.

Despite slight differences in average capsule size both within and between strains, overall, each strain similarly stabilizes to a specific distribution (Figure 4A-C). As with *C. neoformans* strain H99, the most significant differences to capsule distribution were observed after macrophage ingestion (Figure 4D).

### Capsule Skew Benefits ROS Survival and Ingestion Defense

We hypothesized that the reason *C. neoformans* capsule size distributions are so stable is that there was some advantage to this specific capsule distribution. One obvious aspect in which the distribution of capsule size can greatly affect *C. neoformans* population viability is protection from ROS stress, which is produced by immune phagocytic cells during phagosome oxidative burst. It was show that, on a population level, higher average capsule sizes are associated with higher average ROS survival^7^. To investigate this association more thoroughly at the individual rather than populational level, we compared total capsule radius as well as the ratio of capsule to cell body radius to yeast survival post H_2_O_2_ treatment. We found that, as anticipated from these earlier studies, a population of yeast with a larger capsule size are more likely to survive H_2_O_2_ treatment and there is a plateau at which increasing capsule size offered diminishing returns. This plateau shifted depending on the concentration of H_2_O_2_ (Figure 3B). The right-hand tail of capsule sizes ranges far into the protective region.

**Figure 3:**
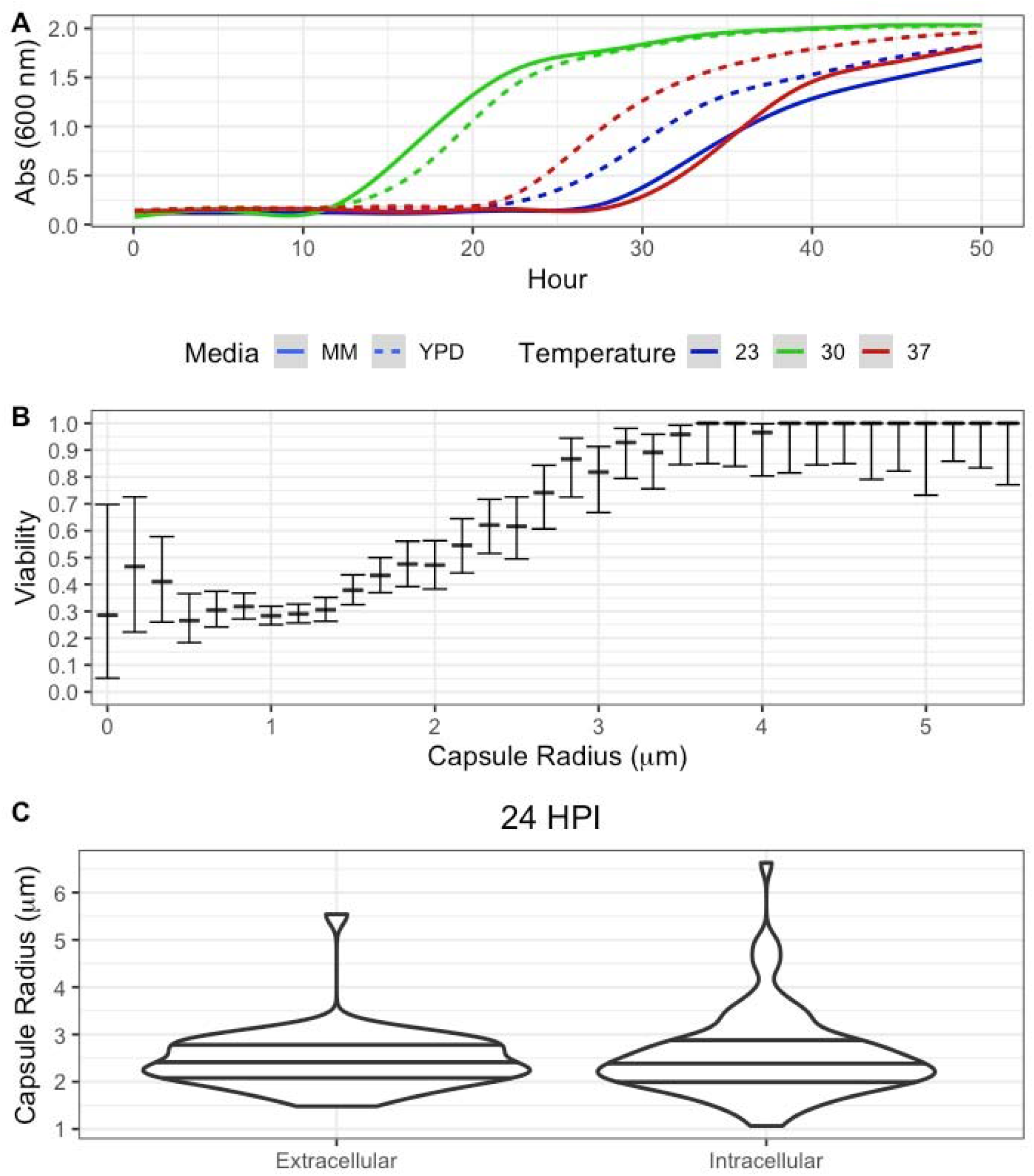
*C. neoformans* strain H99 growth and capsule dynamics in response to cellular stresses. **A.** Growth curves of *C. neoformans* in YPD after preculturing in YPD or MM at environmental (23 °C), optimal (30 °C), and human body (37 °C) temperature. **B**. Viability of *C. neoformans* cultures according to capsule size (rounded to the nearest pixel length) after treatment with 2 mM H_2_O_2_. **C**. Capsule radii of *C. neoformans* 24 h after ingestion by BMDMs. Violins summarize biological triplicate with n = 46 and 85.

**Figure 4:**
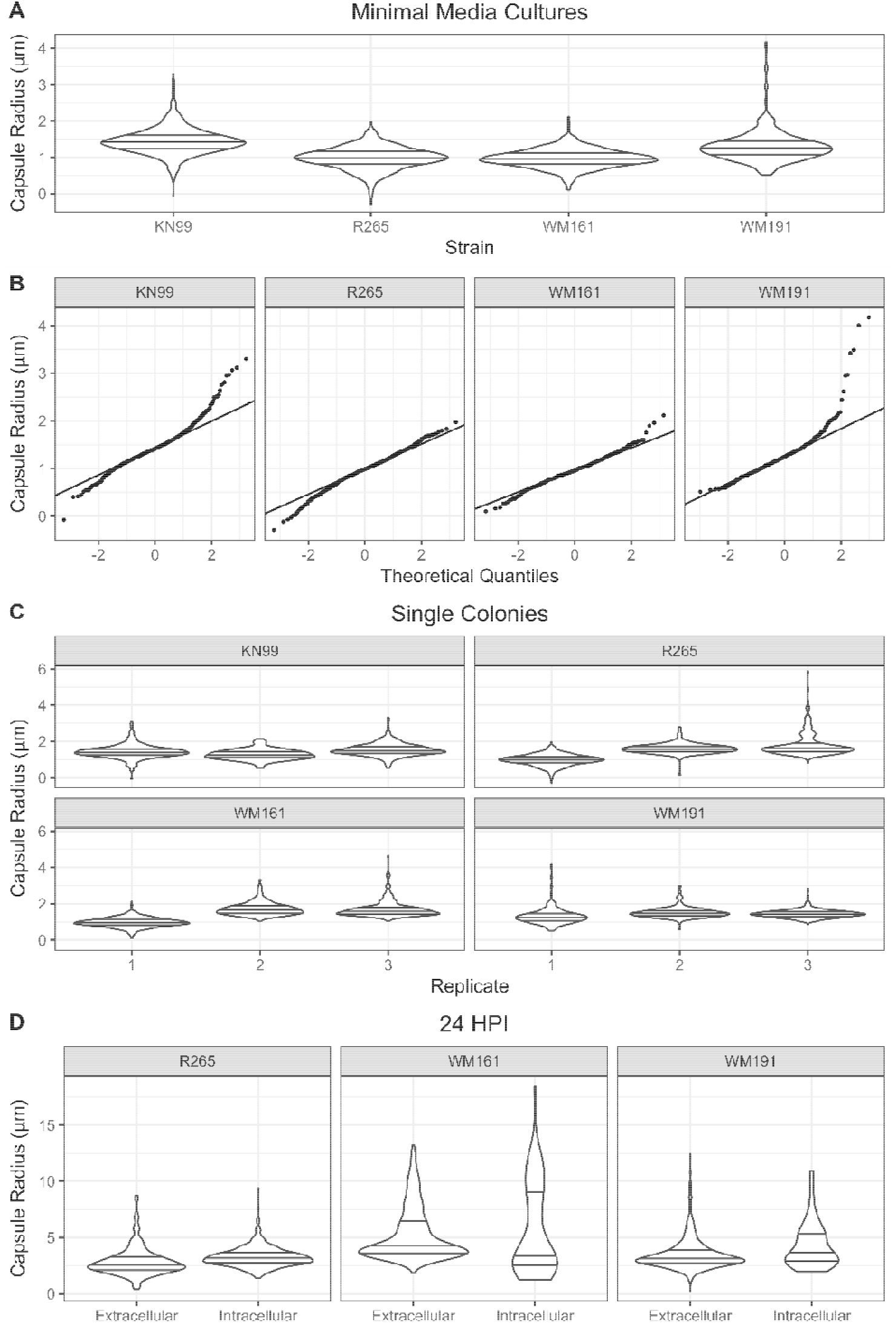
Replicated measurements of additional species: C. neoformans strain KN99 and C. gattii strains WM179, R265, and WM161. **A.** Capsule radius distributions of Cryptococcus strains cultured 7 d in MM. n > 790. **B**. QQ plots of capsule radius distributions of Cryptococcus strains cultured 7 d in MM. **C**. Capsule radiu distributions of Cryptococcus strains cultured from single colonies 7 d in MM. n > 122 **D**. Capsule radiu distributions of Cryptococcus strains 24 HPI of BMDMs. n > 30.

Phagocytes generally struggle with ingesting particles greater than 10 μm and, though uncharacterized, there is likely an upper limit to ingestion size somewhere below the actual size of the phagocyte itself. Indeed, based on an average volume of 4990 μm^3^ for alveolar macrophages, an individual yeast would only need a radius of 10.7 μm to meet this hypothetical upper limit^14^. We found that the heavy right-hand tail in capsule size distribution includes a subset of yeast unlikely to be ingested by alveolar macrophages due to size. Given that amoeba are major environmental predators of Cryptococcus spp. With an incredible range of cell sizes, some much larger than macrophages, this may help explain why the distribution skews greater than what is necessary to survive ROS concentrations *C. neoformans* is likely to encounter in a phagosome^15,16^. However, if larger capsule sizes are guaranteed to better protection and survival, we would instead expect a left-hand skewed population where all yeast are encouraged to produce the largest capsule possible, suggesting there are also disadvantages to large capsules.

### Cryptococcus neoformans Replication Speed Decreases After Capsule Induction at 37 °C

To investigate whether the energy investment required for capsule induction modulated the growth rate of *C. neoformans*, we measured the growth rates of large capsule *C. neoformans* populations compared to small capsule populations after both populations were resuspended in YPD. At optimal growth temperature (30 °C) we found little to no difference in the growth rate of the two conditions. However, room temperature (23 °C) is a more accurate estimate of *C. neoformans* natural environment, at which capsule induced cultures are slower to recover. We observed the same phenotype at human body temperature (37 °C) (Figure 3A).

Interestingly, the energy requirement at a population level shrinks with increased average capsule size, as the number of yeast that can fit in the physical space they are growing in outpaces the energy requirement for the larger capsules (Figure 5). Taken together, these data suggest there is likely an optimal balance between population size and capsule size to increase viability and retain high numbers of yeast while reducing overall energy requirements to match what is available in the environment.

**Figure 5:**
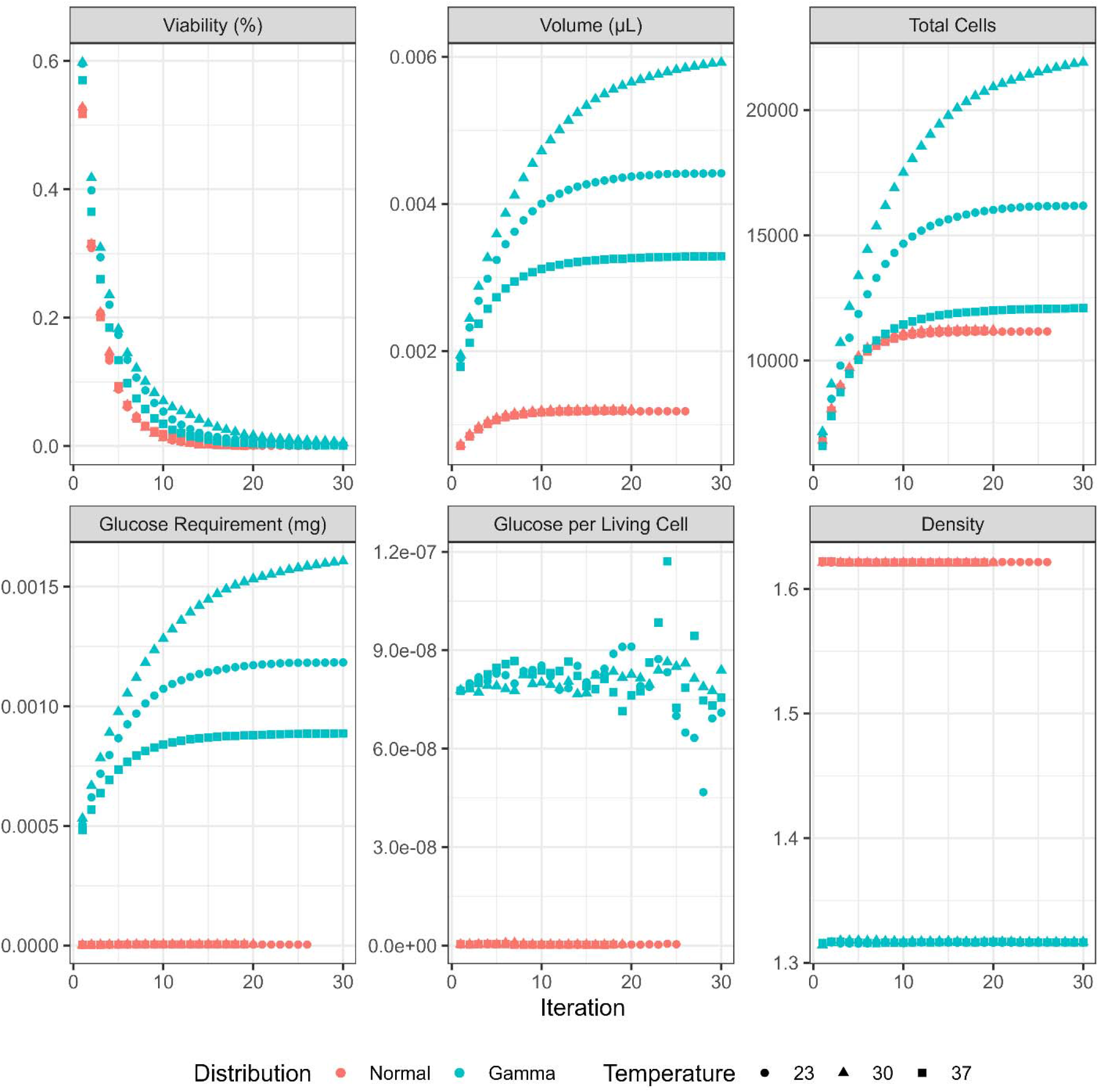
Modeled *C. neoformans* population dynamics at various temperatures under Normal and Gamma distributions. Populations of 5000 initial cells were estimated with capsule sizes of mean 0.839 µm with a standard deviation of 0.219, and cell bodies of mean 2 µm with standard deviation of 0.5. Populations were followed for 30 simulated generations, though the normal distribution completely died off at generation 21.

### A Model for Energy Investiture vs Capsule Benefits

We hypothesized that the observed distributions in capsule sizes resulted from a betting strategy in which the benefit of continually growing a protective capsule was eventually outweighed by the energy, space, and population requirements of generating and maintaining it (Table I). We first modeled the energy required to generate a capsule of a given size according to what was observed using estimates of capsule GXM density, GXM enthalpy, and GXM molecular weight. We were then able to simulate populations of *C. neoformans* under normal and gamma capsule distributions of similar mean size and variance (Figure 5).

As expected, we found that a gamma distribution reflects what is observed in minimal media conditions and confers a survival advantage over a normal distribution when faced with ROS. At all three temperatures the normally distributed population is completely eradicated within 30 generations while the gamma distribution persists. However, modelling selective pressure on a normal distribution did not produce the gamma distribution observed under minimal media conditions among the surviving population, suggesting a change in capsule distribution is actively generated rather than passively emerges from survivor bias, and that additional factors are involved in these capsule dynamics changes.

### Macrophage Ingestion of Yeast Changes Capsule Dynamics

We hypothesized that the distribution of capsule sizes of *C. neoformans* after ingestion by macrophages would resemble the distribution after growth in MM, as both constitute nutritional stress conditions. To investigate these capsule dynamics, we measured capsule sizes of *C. neoformans* every 24 h after ingestion by macrophages. We found no differences in capsule size between ingested or extracellular *C. neoformans*, though both populations were significantly larger than capsules of *C. neoformans* cultured in YPD (Figure 3C). Surprisingly, we found that capsule sizes of macrophage ingested *C. neoformans* were larger on average compared to those cultured in minimal media. It is difficult to parse whether the yeast established a different distribution of capsule sizes when ingested, or whether ingestion and predation by macrophages selects for a certain distribution and we are witnessing survivor bias.

Lysing macrophages to recover internalized yeast produced irregular glob shaped clusters where the cryptococcal cells were encased in a material that held them together rather than appearing as distinct individual cells and capsules. Our initial hypothesis that these zones were cell debris from lysed macrophages was disproven when observing the capsule via IF, noting that capsule specific antibody signal was dispersed throughout the exclusion zone in an irregular, net-like pattern (Figure 6). This material did not diffuse into the lysis solution and had the appearance of a polysaccharide-like component that was produced in the phagosome, which appears to be antigenically different than capsular polysaccharide given mAb staining patterns. Capsule antibody staining patterns were radically altered after the initial 24 h infection period. Massive exclusion zones in India Ink slides began appearing for *C. neoformans* samples opsonized with mAb 13F1, a murine IgM. We also noted that yeast which displayed this net staining pattern qualitatively had lower overall staining intensity than their non-net counterparts, implying either fewer or different monomeric subunits (Figure 6). Interestingly, the ability to overlap with neighboring capsules seems unique to these net capsules. We suspect the net like yeast capsules are important for *C. neoformans* biofilm formation, allowing multiple capsules to adhere to or overlap one another, which could provide additional protection in the phagosome.

**Figure 6:**
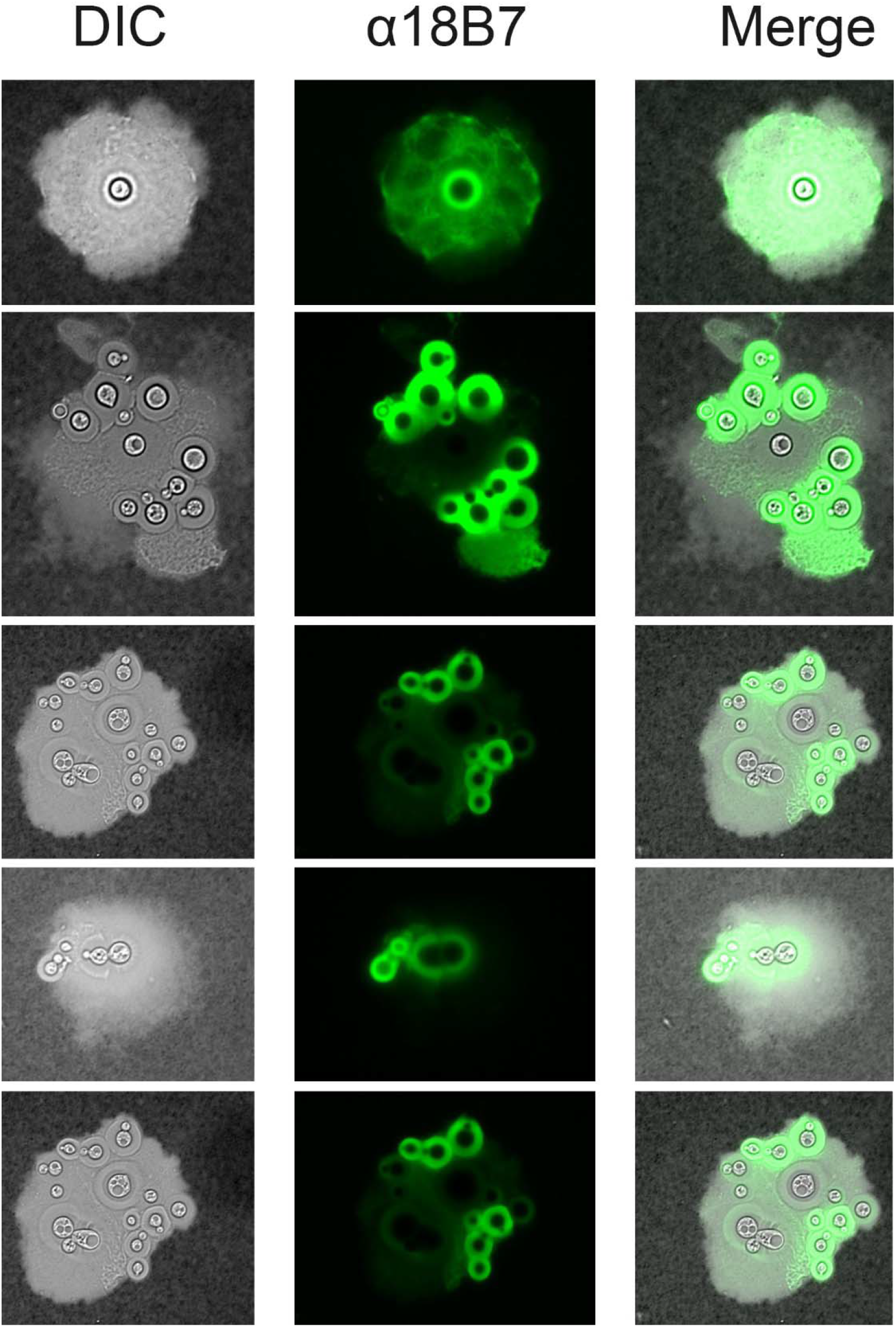
Examples of irregular capsule shapes 24 h post-ingestion by murine macrophages. Columns indicate DIC illumination, α18B7-GFP visualization (Ex488 / Em520), and combined views. Antibody incubation was performed after recovering the ingested yeast from macrophages to avoid inhibiting capsule growth and restructuring.

## Discussion

The unexpected stability in capsule distribution of independent *C. neoformans* populations, regardless of initial condition, is a fascinating observation. We were able to determine that the observed distributions of capsule sizes offer strong benefits in terms of resisting ROS stress at levels higher than expected in a macrophage or amoeba phagolysosome. Hence, we posit the existence of ‘capsulestat’ system of capsule regulation in C. neoformans populations, which produces a consistent distribution of capsule sizes despite significant perturbations such as the addition of fresh media. The term capsulestat is inspired by instruments such as thermostat and rheostat, which control temperature and electric current, respectively. However, rather than a mechanical device the cryptococcal capsulestat is an emergent property from a *C. neoformans* population in which capsule size is determined randomly within the constraints of a bet hedging mechanism and where larger capsule confer more protection at the price of greater metabolic cost.

We first determined that, when grown in glucose rich YPD media, *C. neoformans* defaults to a relatively normal distribution of smaller capsule sizes (approximately 1-1.5 µm). This radius corresponds to a previously observed region of the capsule which is more densely packed with GXM subunit than the rest of the capsule. However, upon capsule induction via nutrient starvation for 5 d, this distribution shifted towards a larger average sized capsules and showed a heavy right-hand tail resulting from the appearance of a population of cells with enlarged capsules. We first hypothesized that this skew was due to the yeast population being divided into fast and slow capsule inducers and that, with time, the distribution would resettle to normality with a higher average capsule radius. However, after allowing cultures to remain in capsule inducing media for several months, these distributions were established early on and never returned to normality. The extreme stability of these capsule size distributions even when supplemented with higher glucose concentrations implies an inherent advantage or selective pressure to establish this specific distribution.

We also observed stochastic dynamics to capsule size when using a scanning window ordinal analysis and that capsule size was not inherited from the progenitor cell, suggesting that capsule radii are randomly chosen at an individual level. We did observe forbidden ordinals in six conditions, however we cannot reject the null hypothesis based on the low samples size of these conditions (<350) ^17^. Our hypothesis is this distribution represents a bet-hedging strategy in which *C. neoformans* must trade energy investment for the protection of a larger capsule. At some point it must become economically nonviable for *C. neoformans* to generate a larger capsule. Producing a larger capsule incurs a replicative cost since capsular enlargement is associated with slower replication rate^6^. However, it may be unnecessary for every yeast to generate the largest capsule possible as smaller capsuled yeast could benefit from buffering effects of larger capsule neighbors, especially in a biofilm.

We considered the benefits of generating a capsule in the context of various cellular stresses that *C. neoformans* is likely to encounter during mammalian infection: temperature, ROS, and ingestion by macrophages. Capsule induction did not have a significant effect on *C. neoformans* replication rate or viability when grown at 30 ^°C^. However, at both 23 ^°C^ and 37 ^°C^ capsule induced populations of *C. neoformans* had a reduced replication rate compared to non-induced populations. We find it likely that capsule induction results in slower population recovery overall, but *C. neoformans* can overcome this reduction at optimal growth temperatures. In certain situations, it could be beneficial for individual microbes in a population to delay initiating a survival function, relying on their neighbors, and saving energy for themselves. It is also possible that *C. neoformans* uses its capsule as a resource depot, storing sugar for future famine conditions. Although capsule recycling for nutrition has not been unequivocally demonstrated in *C. neoformans*, starvation conditions trigger capsular remodeling consistent with the notion that the capsule responds to metabolic stress^18^. We hypothesized that the skewed distribution of capsule sizes in capsule inducing media could result from yeasts delaying their own capsule generation to benefit from their neighbor’s response and that overall capsule radius would diminish over time in unrefreshed conditions as yeast were forced to cannibalize their own capsule for energy.

In fact, the only condition in which we were able to change the distribution of *Cryptococcus* spp. capsule sizes was post-macrophage ingestion. Both intracellular and extracellular yeast recovered after 24 h incubation with macrophages and displayed altered distributions of capsule sizes. All three C. gattii strains exhibited larger capsules compared to C. neoformans after incubation with macrophages. However, it is difficult to know whether the yeast themselves established these new distributions or whether the new distribution is a result of survivor bias, as we were only able to measure yeasts which survived the 24 h infection. Given that our modeling did not observe a shift from normal to gamma distribution under ROS survivor bias, we believe that the observed and modeled distributions of yeast cultures are the result of noise from stochastic variables in the capsule generating molecular system whereas the distributions observed post-ingestion are due to selection by and response to the phagolysosomal environment.

We were only able to characterize an advantage to capsule growth in the context of consistent stress against which a capsule would be effective. Under standard conditions, a normal distribution of capsule sizes observed under YPD media is optimal as it retains the same population size and viability as the increased gamma distribution but with lower glucose and volume requirements. Thus, it remains perplexing why a *C. neoformans* population would induce capsule in minimal media where the main stress is lack of glucose. Perhaps the competition between individual yeasts is a stronger stimulus than populational survival, with individual yeasts racing to store precious glucose as a capsule which also physically pushes away competitors. Perhaps capsule induction can fulfill different roles depending on the situation: a fence to push competitors away during scarcity and a scaffold to pool defenses during stress.

*C. neoformans* capsular dynamics appear to be stochastic which would imply that small differences in capsule assembly for individual cells translate to comparable distributions. The mechanism for capsule growth remains poorly understood but sufficient information is available such that one can conceive a system in which, for a given cell, the final size is a combined result of multiple processes, resulting in a stochastic process. Capsular polysaccharide synthesis begins intracellularly in vesicles that deliver capsular material to the exterior of the cell. *C. neoformans* capsule growth correlates with mitochondrial activity, the production of reactive oxygen species, and depends on numerous additional variables such as metabolism, pH, extracellular vesicle dynamics, and magnesium^2,19–22^. Given that capsule size is regulated at the polymer level and controlled by several signal transduction cascades such as the cAMP/Protein Kinase A (PKA) Pathway, High-Osmolarity Glycerol (HOG) Pathway, Calcium–Calcineurin Signaling, Cell Wall Integrity (CWI) Pathway, RAS1 and RAS2 Pathways, and Iron Regulatory Pathway and that these pathways are noisy and interact, it is not difficult to imagine how each individual cell could have a slightly different capsule size setpoint at any one time that would result in a normal capsule size distribution^23–26^. Experimental evidence for this insights can be found in the temporal growth kinetics of capsules from individual cells, which show great cell-to-cell individual variation ^27^. We propose that in any given culture condition these variables interact to produce a random solution for each cell that, in aggregate, translates to a stable distribution of capsule sizes in the overall population.

When capsule growth and remodeling was not restricted by 18B7 binding, we were able to observe completely different capsule dynamics post macrophage ingestion^28^. We suspect that the distribution of capsules we have observed is a sort of baseline established through stochastic noise in capsule growth related cellular pathway signaling while capsules post-ingestion are heavily modified due to interaction with and response to the phagolysosomal environment.

In summary, the distribution of capsule size across pathogenic *Cryptococcus* spp., strains, and growth conditions manifest remarkable stability. We interpret this stability as providing the population with a bet-hedging strategy whereby replicative fitness is traded for increased fitness from the protection of an enlarged capsule and where the capsule size of individual cells is set by the confluence of numerous, small, chance-dependent influences resulting in stochastic dynamics. Since *Cryptococcus* spp. cells in both environmental and infected host niches cannot anticipate the stresses that they will face from changes in physical and nutritional conditions and from phagocytic cells, and that the capsule is a major protective asset, adopting a capsule size distribution that varies stochastically enhances population survivability through a bet hedging strategy.

---

**Supplemental Figure 1:**
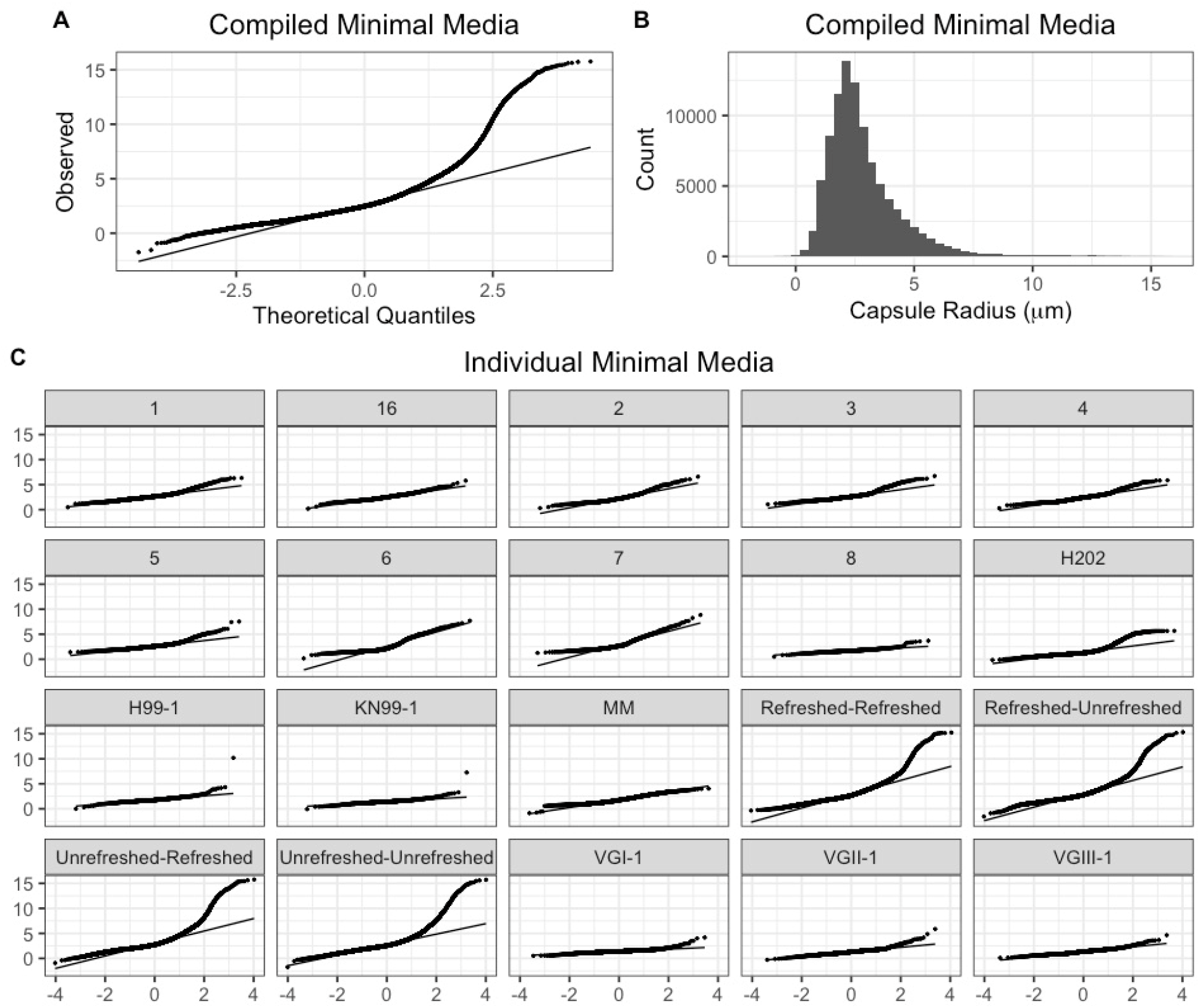
Capsule radius distributions of every experiment and sample in capsule inducing conditions. **A**. QQ plot of capsule distributions for all experiments pooled together. We observe the same right-hand skew as in the initial observation. **B**. Histogram of capsule radii for all experiments pooled together. **C**. Individual QQ plots for each sample and experiment in capsule inducing conditions. While the severity of the right hand skew differs between experiments, the skew is present across each condition.

**Supplemental Table I.** Bond energies in GXM Serotype D. Molecular bond energies estimated in one subunit of GXM minus one molecule H2O which would be lost during polymerization.

| Bond Type | Energy per Bond (kJ / mol) | Quantity in GXM | Total |
| --- | --- | --- | --- |
| C-C | 346 | 21 | 7,266 |
| C-O | 358 | 33 | 11,814 |
| C=O | 799 | 1 | 799 |
| C-H | 413 | 26 | 10,738 |
| O-H | 463 | 14 | 6,482 |
|  |  |  | **37,099** |

